# SARS-CoV-2 S protein ACE2 interaction reveals novel allosteric targets

**DOI:** 10.1101/2020.10.13.337212

**Authors:** Palur Raghuvamsi, Nikhil Tulsian, Firdaus Samsudin, Xinlei Qian, Kiren Purushotorman, Gu Yue, Mary Kozma, Julien Lescar, Peter Bond, Paul MacAry, Ganesh Anand

## Abstract

The Spike (S) protein is the main handle for SARS-CoV-2 to enter host cells through surface ACE2 receptors. How ACE2 binding activates proteolysis of S protein is unknown. Here, we have mapped the S:ACE2 interface and uncovered long-range allosteric propagation of ACE2 binding to sites critical for viral host entry. Unexpectedly, ACE2 binding enhances dynamics at a distal S1/S2 cleavage site and flanking protease docking site ~27 Å away while dampening dynamics of the stalk hinge (central helix and heptad repeat) regions ~ 130 Å away. This highlights that the stalk and proteolysis sites of the S protein are dynamic hotspots in the pre-fusion state. Our findings provide a mechanistic basis for S:ACE2 complex formation, critical for proteolytic processing and viral-host membrane fusion and highlight protease docking sites flanking the S1/S2 cleavage site, fusion peptide and heptad repeat 1 (HR1) as allosterically exposed cryptic hotspots for potential therapeutic development.

**One Sentence Summary:** SARS-CoV-2 spike protein binding to receptor ACE2 allosterically enhances furin proteolysis at distal S1/S2 cleavage sites

---

The COVID-19 pandemic caused by the SARS-CoV-2 virus has sparked extensive efforts to map molecular details of its life cycle to drive vaccine and therapeutic discovery.(*1*) SARS-CoV-2 belongs to the family of *Coronaviridae* which includes other human pathogens including common cold causing viruses (hCoV-OC43, HKU and 229E), SARS and MERS-CoV.(*2–5*) SARS-CoV-2 has a ~30 Kbp long positive RNA genome with 14 open reading frames, encoding 4 structural proteins: Spike (S) protein, membrane (M) protein, envelope (E) protein and nucleoprotein; 16 non-structural proteins and 9 accessory proteins.(*6–8*) An intact SARS-CoV-2 virion consists of a nucleocapsid core composed of nucleoprotein packaged genomic RNA encapsidated as a lipid-protein envelope forming a spherical structure of diameter ~100 nm.(*9*) The viral envelope is decorated with S, M and E proteins.(*9*) The S protein is a club-shaped homotrimeric class I viral fusion protein that has distinctive ‘head’ and ‘stalk’ regions (Fig. 1A).

**Figure 1:**
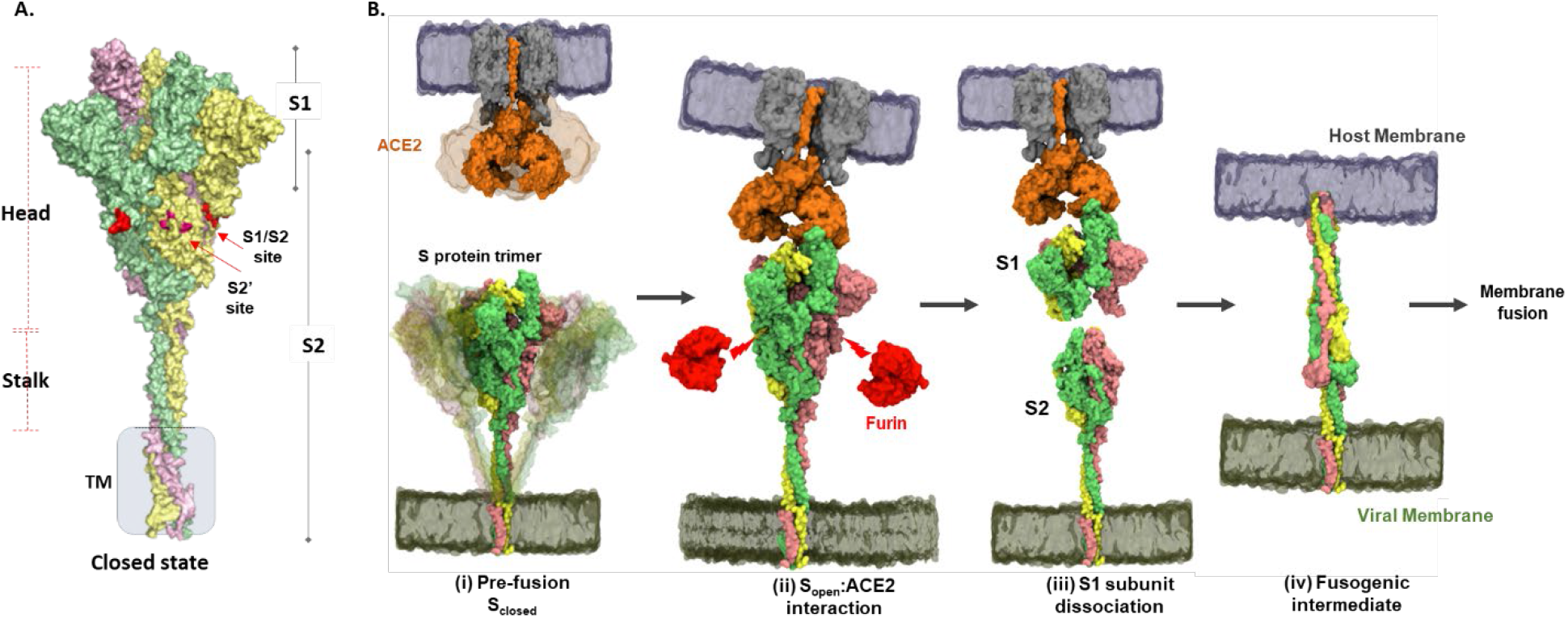
Structure and domain organization of trimeric S protein showing steps in the virushost entry initiated by S recognition and binding to ACE2 receptor. **A.** Prefusion S protein trimer in closed conformational state, with monomers shown in yellow, green and pink. S protein construct (1-1245) used in this study showing head, stalk and transmembrane (TM) segments as generated by integrative modeling. The S1/S2 and S2’ cleavage sites are in red. Proteolytic processing (Furin) of S protein generates S1 and S2 subunits. **B**. Schematic of viral entry into host cell mediated by S:ACE2 interactions as previously outlined(*28*): (i) Intrinsic dynamics of pre-fusion S protein trimer decorating SARS-CoV-2 and host ACE2 dimeric structure showing sweeping motions of S protein and ACE2 to facilitate S:ACE2 recognition. (ii) In the open conformation (S_open_), RBD adopts an ‘up’ orientation to recognize and bind the host membrane-bound ACE2 receptor (PDB: 1R42). ACE2 binding induces conformational changes promoting Furin (red) proteolysis at the S1/S2 cleavage site (red arrows, leading to dissociation of S1 and S2 subunits, mechanism of which is unknown. (iii) The residual ACE2-bound S1 subunit stably bound to ACE2 and S2 subunits dissociate (iv) Conformational changes in the separated S2 subunit promote formation of an extended helical fusogenic intermediate (PDB ID: 6M3W),(*17*) for fusion into the host cell membrane, membrane fusion and viral entry into the host cell. (*11*)

A characteristic feature of SARS-CoV-2 is proteolysis of Pre-fusion S protein by host proteases into S1 and S2 subunits. The S1 subunit comprises an N-terminal domain (NTD) and a receptor binding domain (RBD) that interacts with the host receptor Angiotensin converting enzyme-2 (ACE2)(*10, 11*) to initiate viral entry into the host.(*12*) Cryo-electron tomography (cryo-ET) has been used to capture the distribution and organization of trimeric S protein on the intact virion,(*9*) revealing that 25 ± 9 S protein trimers decorate a single virion with a small percentage (3%) of embedded S proteins in a post-fusion state adopting an extended helical conformation. The first virus-host interaction is mediated by the viral S protein with the host ACE2 receptor.(*10*) Binding to ACE2 primes the S protein for proteolysis at S1/S2 cleavage site into individual S1 and S2 subunits.(*13, 14*) The S2 subunit is divided into six constituent domains harboring the membrane fusion machinery of the virus. These comprise the fusion peptide (FP), heptad repeat (HR1), heptad repeat 2 (HR2), connector domain (CD), transmembrane domain (TM), and cytoplasmic tail (CT).(*15, 16*) Extensive structural studies (*9, 15, 17, 18*) have captured S proteins of coronaviruses in distinct open-(PDB:6VXX)(*15*) and closed-(PDB:6VYB)(*15*) conformational states with regards the RBD, as well as the ectodomain orientation in the pre- and post-fusion states, thereby revealing a high intrinsic metastability of the S protein. The S2 subunit promotes membrane fusion and viral entry (Fig. 1B).

Despite extensive cryo-EM studies, how ACE2 binding at the RBD domain primes enhanced proteolytic processing at the S1/S2 site is entirely unknown. Amide hydrogen/deuterium exchange mass spectrometry (HDXMS) is a powerful complementary tool for both virus dynamics (*19*) and mapping protein-protein interactions.(*20*) Here, we describe the dynamics of free S protein, the S:ACE2 complex and describe ACE2 binding-induced allosteric conformational changes across the distal regions of S protein, particularly at the stalk and protease docking sites flanking the S1/S2 cleavage sites. These distal ‘hotspots’ are critical for the first step of SARS-CoV-2 infection and represent novel targets for therapeutic intervention.

## Results and Discussion

### Localizing subunit specific dynamics and domain motions of S protein trimer

Structural snapshots of the ACE2 binding to the SARS-CoV-2 S protein interface have been obtained with the RBD alone.(*10, 16, 21–23*) In this study, we have mapped this interface for the S protein construct (1-1208) with mutations at the S1/S2 cleavage site (PRRAS to PGSAS) and proline substitution at 986-987,(*16*) to block proteolysis during expression and purification (Fig. S1A). The S protein and isolated RBD constructs showed high affinity binding to ACE2 (Fig. S1B). We measured dynamics of a trimer of this near-full length S protein by amide hydrogen-deuterium exchange mass spectrometry (HDXMS). Pepsin proteolysis generated 321 peptides with high signal to noise ratio, accounting for ~87% of the entire S protein (Fig. S2). Glycosylation of at least 22 sites have been predicted on S protein.(*24*) Average deuterium exchange at these reporter peptides was monitored for comparative deuterium exchange analysis of S protein, ACE2 receptor and S:ACE2 complex, along with a specific ACE2 complex with the isolated RBD. While glycosylation is an important posttranslational modification, our HDXMS study has measured deuterium exchange of non-glycosylated segments of S protein alone. Deuterium exchange (t = 1 and 10 min) across all peptides of the free S protein trimer are shown in (Fig. 2). We built an integrative model of the full-length S protein trimer using experimental structures of prefusion S ectodomain in the open conformation (PDB: 6VSB)(*16*) and the HR2 domain from SARS S protein as templates. Mapping the relative deuterium exchange across all peptides onto this S protein model showed the greatest deuterium exchange at the stalk region. (Fig. 2A) This is consistent with earlier studies showing at least 60° sweeping motions of the three identified hinge regions of the stalk.(*18*) This was further verified via all-atom MD simulations of the S protein model embedded in a viral model membrane, which showed significant motions of the S protein ectodomain as a result of the flexible stalk region, (Fig. 2B) as well as large atomic fluctuations around the HR2 domain, compared to the rest of the protein (Fig. S3, Fig. S4).

**Figure 2:**
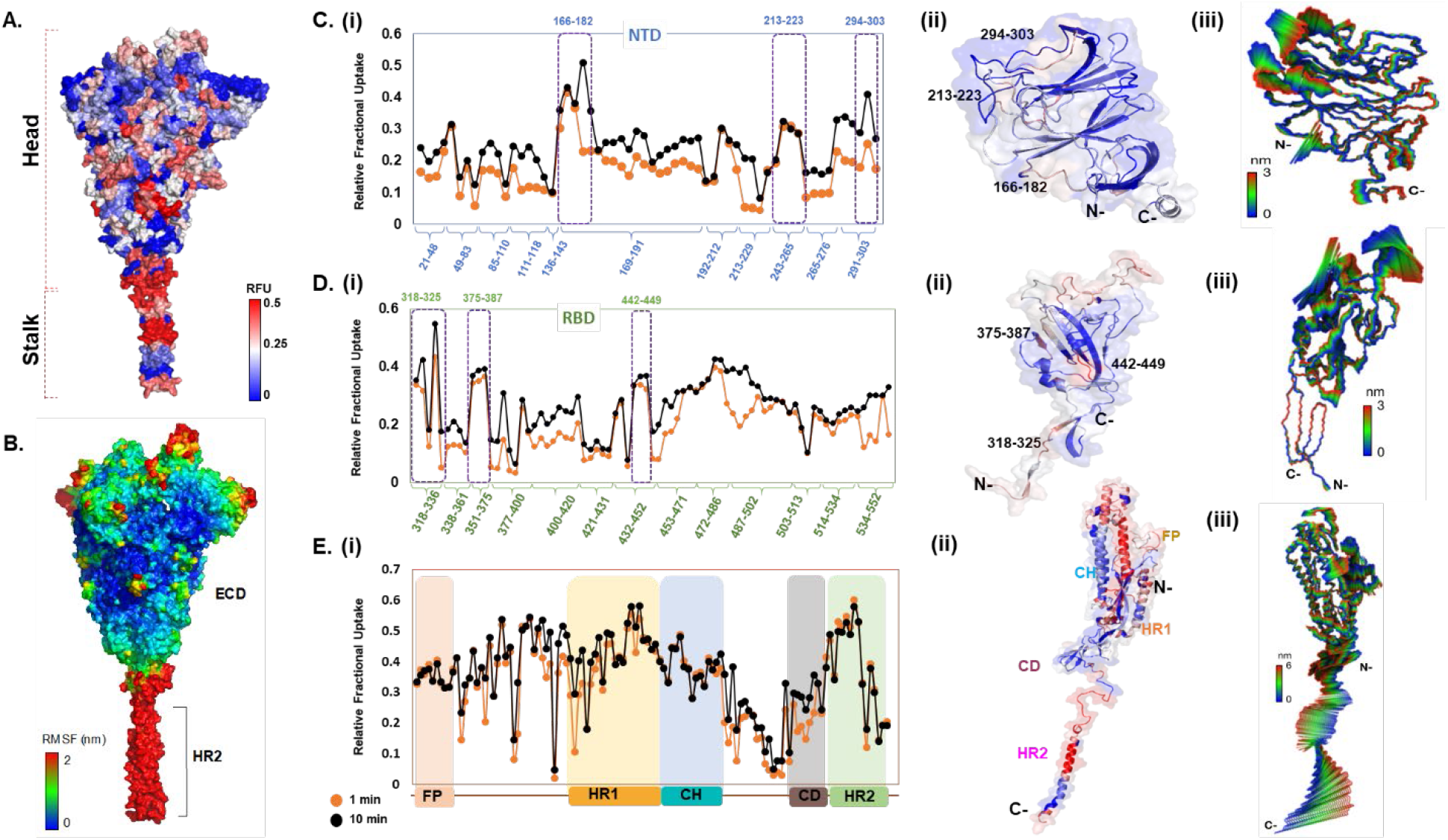
Deuterium exchange heat map and MD simulations reveal domain-specific conformational dynamics of pre-fusion S protein trimer. **A.** Deuterium exchange heat map (shades of blue (low exchange) and red (high exchange)) of S protein (residues 1-1208) at t = 1 min deuterium exchange mapped onto structure of S protein. **B.** Per-residue root mean square fluctuations (RMSF) of the S protein (without TM domain) mapped on to the surface of the S trimer. Deuterium exchange based dynamics across N-terminal domain **(C)**, RBD **(D)**, and the S2 subunit **(E)**. (i) Relative fractional deuterium uptake (RFU) plots of NTD, RBD and the S2 subunit at 1 min (orange) and 10 min (black) deuterium exchange times is shown, with pepsin digest fragments displayed from N to C-terminus (X-axis), (see Fig. S2, Table S1). (ii) Close-up of the structures of NTD (21-303), RBD (318-552) and the S2 subunit (810-1208). Peptides spanning NTD-RBD interaction sites (166-182, 213-223, 294-303, 318-325, 375-387 and 442-449) showing relatively high deuterium exchange at t-=1 min are highlighted. (iii) The first principal motion and RMSF values of backbone atoms on the NTD, RBD and the S2 subunits. Residues with high RMSF are labelled. Different domains (FP, HR1, CH, CD, HR2) showing domain-specific RFU changes are labeled.

The deuterium exchange heat map showed the highest relative exchange in the S2 subunit (Fig. S3) and helical segments, while peptides spanning the fusion peptide showed relatively lower deuterium exchange. Individually, S1 and S2 subunits showed different intrinsic deuterium exchange kinetics, where the average relative fractional deuterium uptake (RFU) of S1 subunit (~0.25) was lower than the average RFU (~0.35) of S2 subunit (Fig. S3, Table S1). Moreover, peptides connecting the RBD to the remainder of the S protein showed greater deuterium exchange, reflecting its role as a ‘hinge’ facilitating the RBD populating an ensemble of open- and closed-conformational states (red arrow, Fig. 2C). Indeed, in our simulations of the S protein (Fig. 2B), the RBD oriented initially in an ‘up’-conformation exhibited spontaneous motion towards the ‘down’-conformation relative to the hinge region (Fig. 2D, Fig. S4A). Interestingly, a part of the receptor binding motif, specifically residues 476-486, exhibited a higher degree of flexibility based on its average atomic fluctuations (Fig. 2A, 4C), suggesting that binding to ACE2 receptor would be required to stabilize its motion.

The NTD of the S protein showed low overall RFU (~0.2), consistent with its well-structured arrangement of β-sheets connected by loops (Fig. 1B). Importantly, certain regions showed significantly higher deuterium exchange (~0.4), of which two loci (136-143, 243-265) span the dynamic interdomain interactions with the RBD. This is supported by the high per-residue root mean square fluctuations (RMSF) and large principal motions observed for residues 249-259 during simulations (Fig. 2C, Fig. S4C). One locus (291-303) at the C-terminal end of the NTD connecting to the RBD showed high deuterium exchange, indicative of relative motions of the two domains. The RBD (Fig. 1D) showed relatively higher deuterium exchange (RFU ~0.35), with the peptides spanning the hinge-regions (318-336) showing greatest deuterium exchange (~0.6). Peptides spanning residues 351-375 and 432-452 showed significantly increased deuterium uptake, and these correspond to the NTD interdomain interaction sites. Interestingly, loci of the RBD implicated in the interface (453-467, 491-510) with ACE2 showed relative higher exchange.

Overall, the S2 subunit showed relatively higher RFU than the S1 subunit, with each domain exhibiting specific conformational changes (Fig. 1E, Fig. S4). Peptides spanning the region immediately downstream of the S1/S2 cleavage site showed the highest deuterium uptake (0.6), reflecting the rapid dynamics it undergoes for facilitating cleavage of S protein into two subunits. Congruently, our MD simulations revealed the unstructured loop housing the S1/S2 cleavage site (residues 677-689) to be highly dynamic (Figure S4C), with RMSFs reaching >1.0 nm. It is important to note that the S1/S2 cleavage site has been abrogated in the construct of the S protein used in this study to block proteolytic processing into S1 and S2 subunits during expression in host cells. We thus observed lower deuterium uptake (and lower RMSF values) at peptides in the central helix and connector domain, suggesting that these act as the central core of prefusion S, while the peptides spanning hinge-segments and heptad repeats (HR1 and HR2) showed high deuterium uptake and RMSF values, indicative of the inherent metastability of S to adopt prefusion, fusion and post-fusion conformations.

### Domain-specific and global effects of ACE2 binding to the RBD

Comparative HDXMS analysis of the S protein and S:ACE2 complex revealed large-scale effects upon binding of ACE2. The main target for direct interactions was the RBD. We therefore set out to characterize the effects of ACE2 binding with RBD (‘RBD_S_’) present on full S protein (Fig. 4A, 4B) and compared this to an isolated construct of the RBD (‘RBD_isolated_’) (Fig. 3, Fig. S6). Several peptides of the RBD_S_ showed decreased exchange upon complexation with ACE2 (Fig. 3B). These include peptides 340-359, 400-420, 432-452 and 487-502 in the RBD_S_:ACE2 complex (Fig. 4). These sites are consistent with the interface of the SARS-CoV-2 S protein RBD bound to the ACE2 receptor resolved by X-ray crystallography.(*10*) The high-resolution structures showed that the RBD and ACE2 receptor interact via an extensive interface. However, not all peptides at the interface contribute equally to the binding energetics. HDXMS reveals the residues at the core of this interface to be those within peptides spanning residues 340-359, 400-420, 432-452 and 491-510 (Fig. 4A, 4D, Fig. S3). Interestingly, loci showing large-magnitude deuterium exchange correlate to mutational hotspots(*25*).

**Figure 3.**
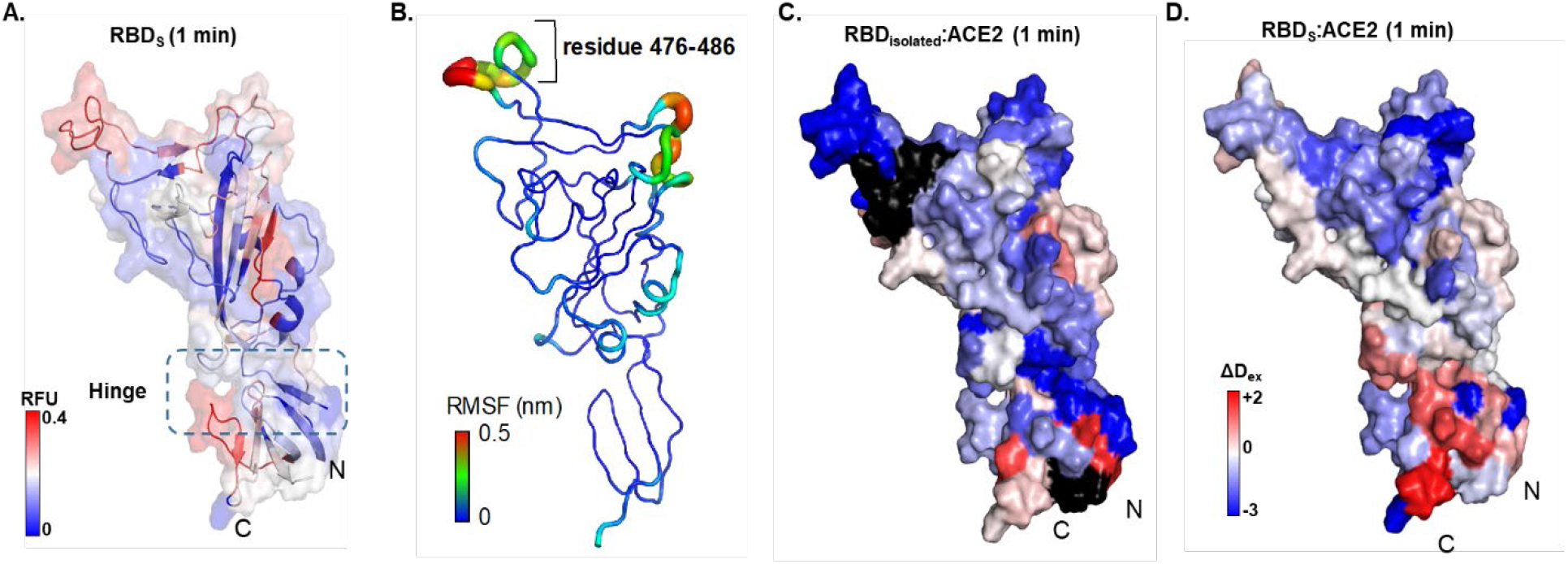
Map of RBD_isolated_:ACE2 interactions. **(A)** Relative fractional deuterium uptake values at t = 1 min for RBD (314-547) of S protein (RBD_S_) mapped on to the structure of RBD extracted from S protein model (see Table S2). High and low exchanging regions are represented as shown in key, and regions with no coverage are shown in black. (**B**) The RMSF values of backbone atoms on the RBD showing residues with high RMSF (476-486), as per key. Differences in deuterium exchanged between RBD_isolated_:ACE2 complex and free RBD_isolated_ **(C)** and RBD_spike_:ACE with free RBD_spike_ **(D)** at 1min of deuterium labelling are mapped on to the structure of RBD. Protection from deuterium uptake and increases in exchange are indicated in blue and red respectively. Regions with no coverage are in black.

A close-up of RBD_isolated_:ACE2 complex also showed decreased exchange in peptides spanning these regions (Fig. 3). However, the magnitude of decrease in exchange was significantly greater in RBD_isolated_ than in RBDS, indicating the higher flexibility in the full-length S trimer than in free RBD. High resolution structures have identified the RBD interface interacting with ACE2 spanning the peptide covering residues 448-501 (Y449, Y453, N487, Y489, G496, T500, G502, Y505, L455, F456, F486, Q493, Q498 and N501) using only the RBD from the S protein.(*23*) Cryo-EM studies have shown that each RBD in the trimeric S protein can adopt an open conformation irrespective of other RBDs, indicating an absence of cooperativity between the three RBDs within a trimer.(*9*) Therefore, we compared the deuterium exchange profiles of RBD_isolated_ with RBDS and observed differences in dynamics imposed by quaternary contacts (Fig. 3A, Fig. 3B). Overall, the loci with high and low deuterium exchange profiles were similar when compared between RBD_isolated_ and RBDS, both at the disordered ACE2 receptor binding region as well as the folded region at the N- and C -termini. In solution RBDS toggles between open- and closed-conformations resulting in an average readout of deuterium exchange measurements.

ACE2 binding to RBD_isolated_ and RBDS resulted in similar effects, where we observed deuterium exchange protection at the peptide regions spanning the known binding interface of RBD. Notably, increased deuterium exchange was observed at the hinge region (Fig. 3C, Fig. S4) indicating allosteric conformational changes, associated with restricting the open- and closed-states interconversion. Therefore, the destabilization/ local unfolding observed at the hinge region as a result of ACE2 binding enables RBD to maintain open conformation. It therefore seems likely that small molecules and biologics targeting the hinge region to lock RBD in the closed state would be of potential high therapeutic value.

### ACE2 binding at the RBD is allosterically propagated to the S1/S2 cleavage site and Heptad Repeat

Unexpectedly, ACE2 binding at the RBD induced large-scale changes in deuterium exchange in distal regions of the S protein. Some of the peptides in the stalk of S protein showed decreased exchange in the S:ACE2 complex (Fig. 4C, 4D). This indicates that ACE2 receptor interactions stabilized the hinge dynamics. Decreased exchange was also seen in the distal sites in the S2 subunit, localized at the fusion peptide locus and central helix (CH). Interestingly, increased exchange was seen in multiple peptides flanking the S1/S2 cleavage site, HR1 domain and critically at the S1/S2 cleavage sites (Fig. 4D). Even though the construct used in this study has the proteolysis site mutated, it still resulted in increased dynamics at this S1/S2 locus. Furthermore, this region exhibited high RMSF values during simulations. (Fig. S4B). These results clearly indicate that ACE2 binding induces allosteric enhancement of dynamics at this locus, providing mechanistic insights into the conformational switch from the pre-fusion to fusogenic intermediate. Differences in deuterium exchange between free S protein and S:ACE2 complex shows stabilization at ACE2 interacting site and local destabilization at peptides juxtaposed to S1/S2 cleavage site (residues 931-938). This suggests that ACE2 binding potentiates peptide of residues 931-938 and other high exchanging regions flanking the S1/S2 cleavage site for enhanced furin protease binding and cleavage. Importantly, these results suggest that the S1/S2 cleavage site is a critical hotspot for S protein dynamic transitions for viral entry into the host, and therefore represents a new target for inhibitory therapeutics against the virus.

**Figure 4:**
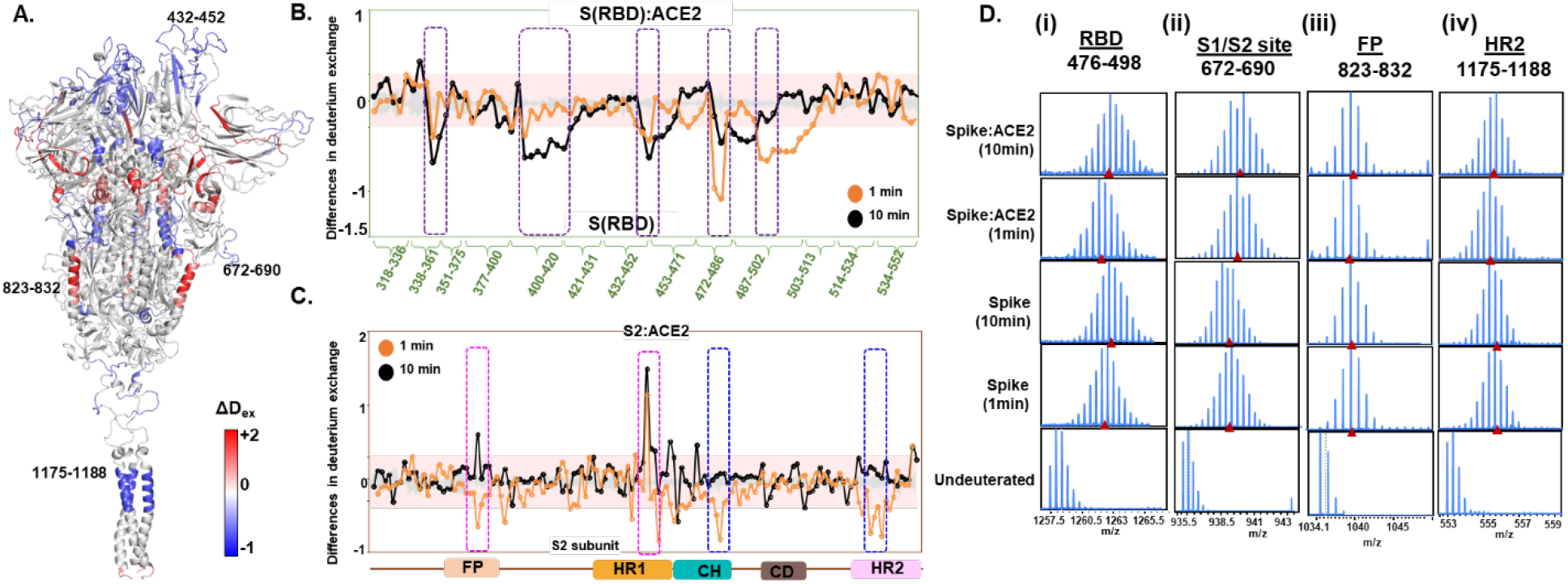
ACE2 interaction induce large scale allosteric changes across S protein. **(A)** Differences in deuterium exchange (ΔD_ex_) (t = 1 min) in S protein upon binding ACE2 showing decreased (blue) and increased (red) deuterium exchange, mapped onto structure of S protein. These differences in deuterium exchange for peptides from **(B)** RBD and S2 subunit **(C)** for pepsin digest fragments (X-axis) are shown. Difference cutoff ±0.3 D is the deuterium exchange significance threshold indicated by pink shaded box with standard error values in gray. Positive differences (>0.3 D) denote increased deuterium exchange and negative differences (<-0.3 D) denote decreased deuterium exchange in S protein bound to ACE2. **(B)** Peptides spanning residues interacting with ACE2 are in purple. **(C)** Peptides spanning fusion peptide (FP) and HR1 are highlighted in pink boxes, while peptides spanning central helix (CH) and heptad repeat 2 (HR2) are in blue. **D.** Stacked mass spectra with isotopic envelopes after deuterium exchange (t = 1, 10 min) for select peptides from (i) RBD (residues 476-498), (ii) S1/S2 cleavage site (residues 672-690), (iii) fusion peptide (residues 823-832) and (iv) HR2 (residues 1175-1188) are shown for the S protein and S:ACE2 complex. Mass spectra of the equivalent undeuterated peptide are shown for reference. The centroid masses are indicated by red arrow-heads.

### Dynamics of ACE2 with RBD and S protein provides insights into viral-entry into host

Considering the indispensable role of ACE2 binding in SARS-CoV-2 infection, it is crucial to assess the effects of S protein and RBD binding on ACE2 dynamics. We therefore mapped the corresponding binding sites of RBD, both isolated and within the Spike, on ACE2. The S:ACE2 complex represents the prefusion pre-cleavage state wherein full-length S protein is bound to the ACE2 receptor (Fig. 1B ii), while the RBD_isolated_:ACE2 complex represents the post-furin cleavage product formed by the S1 subunit and ACE2 (Fig. 1B iii). Previous studies have shown that 14 key amino acids of RBD interact with ACE2, wherein mutations at 6 amino acids resulted in higher binding affinity of SARS-CoV-2.(*26*) SARS-CoV-2 adopted a different binding mode to ACE2 as a superior strategy for infection in comparison to SARS-CoV-1. A crystal structure of RBD_isolated_:ACE2 complex has identified 24 key ACE2 residues, spanning across peptides 16-45, 79-83, 325-330, 350-357 and Arg393.(*27*) While most of these residues are conserved in binding to both SARS-CoV-1 and SARS-CoV-2, Arg393 and residues 325-330 are unique to SARS-CoV-1 interaction.(*25*) Interestingly, we observed increased deuterium exchange at these residues in the S:ACE2 complex compared to ACE2 alone (Fig. S8). Identifying the intrinsic dynamics and allosteric changes due to binding could potentially better inform drug development.

Simulations of the ACE2 dimer complexed with the B^0^AT1 amino acid transporter (PDB: 6M1D)(*12*) in a model epithelial membrane revealed a large motion of the peptidase domain (PD), which recognizes the S protein RBD, with respect to the transmembrane and juxtamembrane domains (Fig. S7). This large motion is reminiscent of the flexible tilting displayed by the S protein ectodomain itself, suggesting that both S protein and ACE2 have adaptable hinges that allow for orientational freedom of the domains involved in recognition. To understand how S protein binding affects ACE2 dynamics, we performed HDXMS experiments of monomeric ACE2 alone, S:ACE2 and RBD:ACE2 complexes (Fig. S7, Fig. S8) and mapped the deuterium exchange values on a deletion construct of ACE2 (PDB: 1R42)(*27*) (Fig. S7, Fig. S8). We observed a reduction in deuterium exchange across both RBDisolate:ACE2 and larger S:ACE2 complexes compared to free ACE2 (Figure S8B and S8C). Differences in deuterium exchange between RBD_isolated_:ACE2 complex and free ACE2 showed that RBD binding stabilizes ACE2 globally, specifically large differences at the binding site (peptides 21-29, 30-39, and 75-92), and also at distal regions (peptides 121-146, 278-292, 575-586) from RBD binding site of ACE2 (Fig. S8D). Cryo-EM studies have shown that a dimeric full length ACE2 receptor can stably bind to one trimer of the S protein.(*12*)

## Conclusions

Here a combination of HDXMS and MD simulations provide a close-up of S protein dynamics in the pre-fusion, ACE2 bound and other associated conformations. Our results reveal the energetics of the S:ACE2 complex interface. ACE2 binding to the isolated RBD and S protein alike lead to binding and stabilization. Interestingly, ACE2 binding to the RBD induces global conformational changes across the entire S protein. Importantly, the stalk region undergoes dampening of conformational motions while causing increased deuterium exchange in the protease sites. Regions highlighting the allosteric propagation of ACE2 binding represent cryptic targets for small molecule inhibitor/antibody development as therapeutics.

## Abbreviations

HDXMS: Hydrogen Deuterium Exchange Mass Spectrometry;
MD: molecular dynamics,
RFU: Relative Fractional deuterium uptake;
RMSF: root mean squared fluctuations;
PCA: Principal Component Analysis;
S: Spike;
UPLC: Ultra Performance Liquid Chromatography;

## Acknowledgements

We thank Dr. Lu Gan, Dept. of Biological Sciences, National University of Singapore for helpful discussions. We thank Protein Production Platform of Nanyang Technological University for their help in making the RBD and ACE2 expression constructs and small-scale protein expression tests. HDXMS experiments were carried out as a fee for service at the Singapore National Laboratory for Mass Spectrometry (SingMass) funded by NRF, Singapore. P.V.R. was supported by research scholarship from National University of Singapore, Singapore. N.K.T. was supported by research grant from Ministry of Education, Singapore awarded to G.S.A. (MOE2017-T2-A40-112). This work was supported by BII of A*STAR. Simulations were performed on the petascale computer cluster ASPIRE-1 at the National Supercomputing Centre of Singapore (NSCC) and the A*STAR Computational Resource Centre (A*CRC).

## Author Contributions

Conceptualization – P.V.R., N.K.T., P.A.M., G.S.A., P.J.B.; Funding acquisition – P.A.M; Investigation – P.V.R., N.K.T., F.S., X.Q., K.P., G.Y., M.M.K.; Methodology – P.V.R., N.K.T., F.S.; Resources – P.V.R., N.K.T., P.A.M., G.S.A., P.J.B.; Supervision & Validation; Visualization – P.V.R., N.K.T., F.S.; Writing – original draft – P.V.R., N.K.T., G.S.A., F.S., P.J.B.; Writing – review & editing – P.V.R., N.K.T., G.S.A., F.S., P.J.B., P.A.M.

## Data and Materials Availability

All data is available in the main text or the supplementary materials. Data, code and materials provided upon request.

## SUPPLEMENTARY MATERIALS

### Materials and Methods

#### Materials

Mass Spectrometry grade acetonitrile, formic acid and water were from Fisher Scientific (Waltham, MA); Deuterium oxide was from Cambridge Isotope Laboratories (Tewksbury, MA). All reagents and chemicals were research grade or higher and obtained from Merck-Sigma-Aldrich (St. Louis, MO).

#### Methods

##### Transient expression and purification of recombinant SARS-CoV2 Spike, RBD and ACE2 receptor

A near full-length Spike (S) protein, excluding transmembrane domain and cytoplasmic tail, of SARS-CoV-2 (1-1208; Wuhan-Hu-1; GenBank: QHD43416.1) was codon optimized for mammalian cell expression and cloned into pTT5 expression vector (National Research Council Canada, NRCC) with a twin strep tag at the C-terminus (Twist Biosciences). This double mutant Spike construct was generated by mutating RRAR (682-685) into GSAS and residues KV (986-987) into PP. A gene encoding SARS-CoV-2-RBD (319-591 of SARS-CoV-2 Spike) (BioBasic) was cloned into the expression vector pHLmMBP-10 (Addgene) following the N-terminal His and mMBP tag. A gene encoding human ACE2 (residues 21-597; GenBank: AB046569.1) fused to a C-terminal Fc tag (Biobasic) was cloned into vector pHL-sec (Addgene) between the signal peptide and c-terminal His tag. SARS-CoV-2-Spike constructs were expressed in HEK293-6E (NRCC) using polyethylenimine (PEI) as the transfection reagent while the isolated RBD (‘RBD_isolated_’) and ACE2 constructs were expressed in Expi293F using the Expi293 System (Thermo Fisher). Culture supernatant was harvested on day 7 for HEK293-6E expression and day 5 for Expi293F expression. Spike proteins were affinity purified using Strep-Tactin^®^XT column (IBA). RBD protein was affinity purified using cOmplete™ His-Tag Purification column (Merck). ACE2 receptor was affinity purified using HiTrap^®^ MabSelect™ SuRe™ column (GE Healthcare). Purified proteins were concentrated and buffer exchanged into PBS using VivaSpin (Sartorius) and the purity was assessed by denaturing polyacrylamide gel electrophoresis (Fig. S1A).

##### Deuterium labelling and quench conditions

Recombinant purified S protein (8 μM), ACE2 receptor (52 μM) and RBD (67 μM) solubilized in phosphate buffer (PBS, pH 7.4) were incubated at 37°C in PBS buffer reconstituted in D_2_O (99.90%) resulting in a final D_2_O concentration of 90%. S:ACE2 and RBD:ACE2 complexes (K_D_ of ~15 nM and ~150 nM, respectively)(*29*) were pre-incubated at 37°C for 30 min in a molar ratio of 1:1 to achieve >90% binding prior to each hydrogen-deuterium exchange reaction. Deuterium labeling was performed for 1 min, 10 min and 100 min for isolated construct of RBD, free ACE2, and RBD_isolated_:ACE2 complex. For isolated S protein and S:ACE2 complex 1 min and 10 min labelling timescales were used. Pre-chilled quench solution 1.5 M GnHCl and 0.25 M Tris(2-carboxyethyl) phosphine-hydrochloride (TCEP-HCl) was added to deuterium exchange reaction mixture to lower the pH_read_ to ~2.5 and lower temperature to ~ 4 °C. Next, the quenched reaction was incubated at 4 °C on ice for 1 min followed by pepsin digestion.

##### Mass Spectrometry and peptide identification

~100 pmol quenched samples were injected onto chilled nanoUPLC HDX sample manager (Waters, Milford, MA). The injected samples were subjected to online digestion using immobilized Waters Enzymate BEH pepsin column (2.1 × 30 mm) in 0.1% formic acid in water at 100 μl/min. Simultaneously, the proteolyzed peptides were trapped in a 2.1 × 5 mm C18 trap (ACQUITY BEH C18 VanGuard Pre-column, 1.7 μm, Waters, Milford, MA). Following pepsin digestion, the proteolyzed peptides were eluted using acetonitrile gradient of 8 to 40 % in 0.1 % formic acid at a flow rate of 40 μl min^-1^ into reverse phase column (ACQUITY UPLC BEH C18 Column, 1.0 × 100 mm, 1.7 μm, Waters) pumped by nanoACQUITY Binary Solvent Manager (Waters, Milford, MA). Electrospray ionization mode was used to ionize peptides sprayed onto SYNAPT G2-Si mass spectrometer (Waters, Milford, MA) acquired in HDMS^E^ mode of detection and measurement. A flow rate of 5 μl/min was used to inject 200 fmol μl^-1^ of [Glu^1^]-fibrinopeptide B ([Glu^1^]-Fib) into mass spectrometer for lockspray correction.

Undeuterated protein samples were used to identify sequences from mass spectra data (in HDMS^E^ mode) using Protein Lynx Global Server (PLGS) v3.0. Peptide identification search was performed against a separate sequence database of each protein sequence along with its respective affinity purification tag sequences. In the PLGS search parameters, i) no specific protease and ii) no variable N-linked glycosylation modification options were selected for sequence identification. The identified peptides were further filtered using a minimum intensity cutoff of 2500 for product and precursor ions, minimum products per amino acids of 0.2 and a precursor ion mass tolerance of <10 ppm using DynamX v.3.0 (Waters, Milford, MA) and tested for pepsin cleavage specificity.(*30*) Peptides independently identified under the specified condition and present in at least in two out of three undeuterated samples were retained for HDXMS analysis. S protein contains 22 variable glycosylation sites(*31*) out of which we identified peptides spanning 12 glycosylation sites in our sample (Fig. S2). For ACE2, we obtained 4 peptides overlapping the glycosylation sites (Fig S7). Relative fractional deuterium uptake (RFU) is the ratio of number of deuterons exchanged to the total number of exchangeable amides of the peptide. Centroid masses of undeuterated reference spectra were subtracted from equivalent spectra of peptides showing deuterium exchange to calculate the average deuterons exchanged with time for each peptide. Deuterium exchange plots, relative deuterium exchange and difference plots were obtained from DynamX 3.0. N-terminus and prolines were excluded for estimation of exchangeable amides per peptide.(*32*) All deuterium exchange experiments were performed in triplicate and reported values are not corrected for deuterium back exchange.

##### Modelling and molecular dynamics (MD) simulations

An integrative model of full-length SARS-CoV-2 S protein was built using Modeller v9.21.(*33*) The cryo-EM structure of pre-fusion S ectodomain in the open conformation (PDB: 6VSB)(*29*) was used as the template for the ectodomain (ECD) with missing loops on the NTD modelled based on SARS S NTD crystal structure (PDB: 5X4S).(*34*) The NMR structure of SARS S HR2 domain (PDB: 2FXP)(*35*) was used as the template for the HR2 domain, while the TM domain was modelled using the NMR structure of the HIV-1 gp-41 TM domain (PDB: 5JYN)(*36*). Ten models were built and subjected to stereochemical assessment using the discreet optimized protein energy (DOPE) score(*37*) and Ramachandran analysis.(*38*) The model with the lowest DOPE score and the smallest number of Ramachandran outliers was chosen. Palmitoylation was added to three cysteine residues (C1236, C1240 and C1243) on the CT domain based on a study showing its importance in SARS S protein function.(*39*) The S protein model was then embedded into a model membrane representing the endoplasmic reticulum-Golgi intermediate compartment (ERGIC),(*40*) where coronaviruses are known to assemble in a bud *form*.(*41, 42*) The ERGIC model membrane was built using CHARMM-GUI Membrane Builder.(*43*) All-atom MD simulation was performed for 200 ns using GROMACS 2018(*44*) and the CHARMM36 force field.(*45*) The systems were solvated with TIP3P water molecules and 0.15 M NaCl salt. Minimization and equilibration were performed following standard CHARMM-GUI protocols.(*46*) Temperature was maintained at 310 K using the Nosé-Hoover thermostat(*47, 48*) and the pressure was maintained at 1 atm using the Parrinello-Rahman barostat.(*49*) Electrostatics were calculated using the smooth particle mesh Ewald (PME) method(*50*) with a real space cutoff of 1.2 nm and the van der Waals were truncated at 1.2 nm with force switch smoothing between 1.0 to 1.2 nm. Constraints were applied to covalent bonds with hydrogen atoms using the LINCS algorithm(*51*) and a 2 fs integration time step was employed. For simulations of the ACE2 receptor, the cryo-EM structure of ACE2-B^0^AT1 complex in the open conformation (PDB: 6M1D)(*52*) was used. The ACE2-B^0^AT1 complex was embedded into a model membrane representing the epithelial cell membrane.(*53, 54)* All-atom MD simulation was performed for 200 ns using the protocols described above. Principal component analysis (PCA) and root means square fluctuation (RMSF) analyses were performed using GROMACS, and simulations were visualized in VMD.(*55*)

## Supplementary Figures

**Figure S1:**
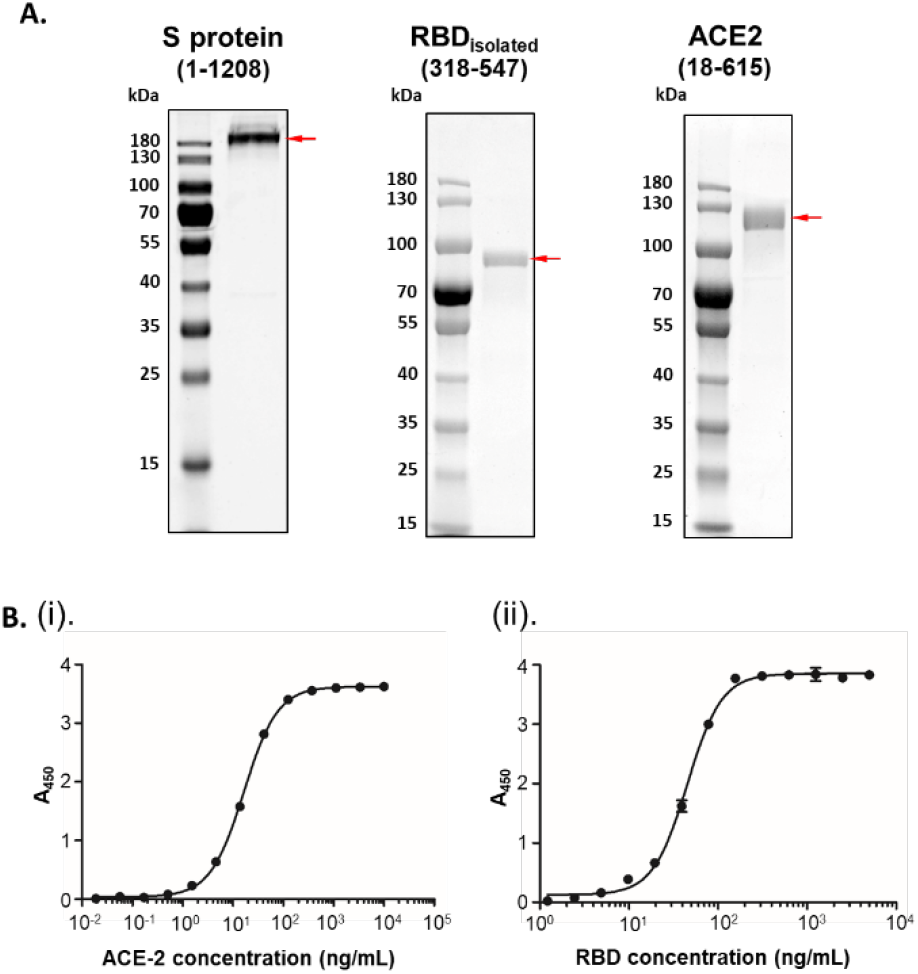
Homogeneity of protein samples. (A) Images of denaturing polyacrylamide gel electrophoresis of purified proteins of the S protein (mutant), isolated RBD and ACE2 are shown, and their molecular sizes are highlighted with red arrow, alongside protein standards. Domain organization is shown for reference. (B) Interactions between ACE2 and RBD represented by the binding curves.

**Figure S2:**
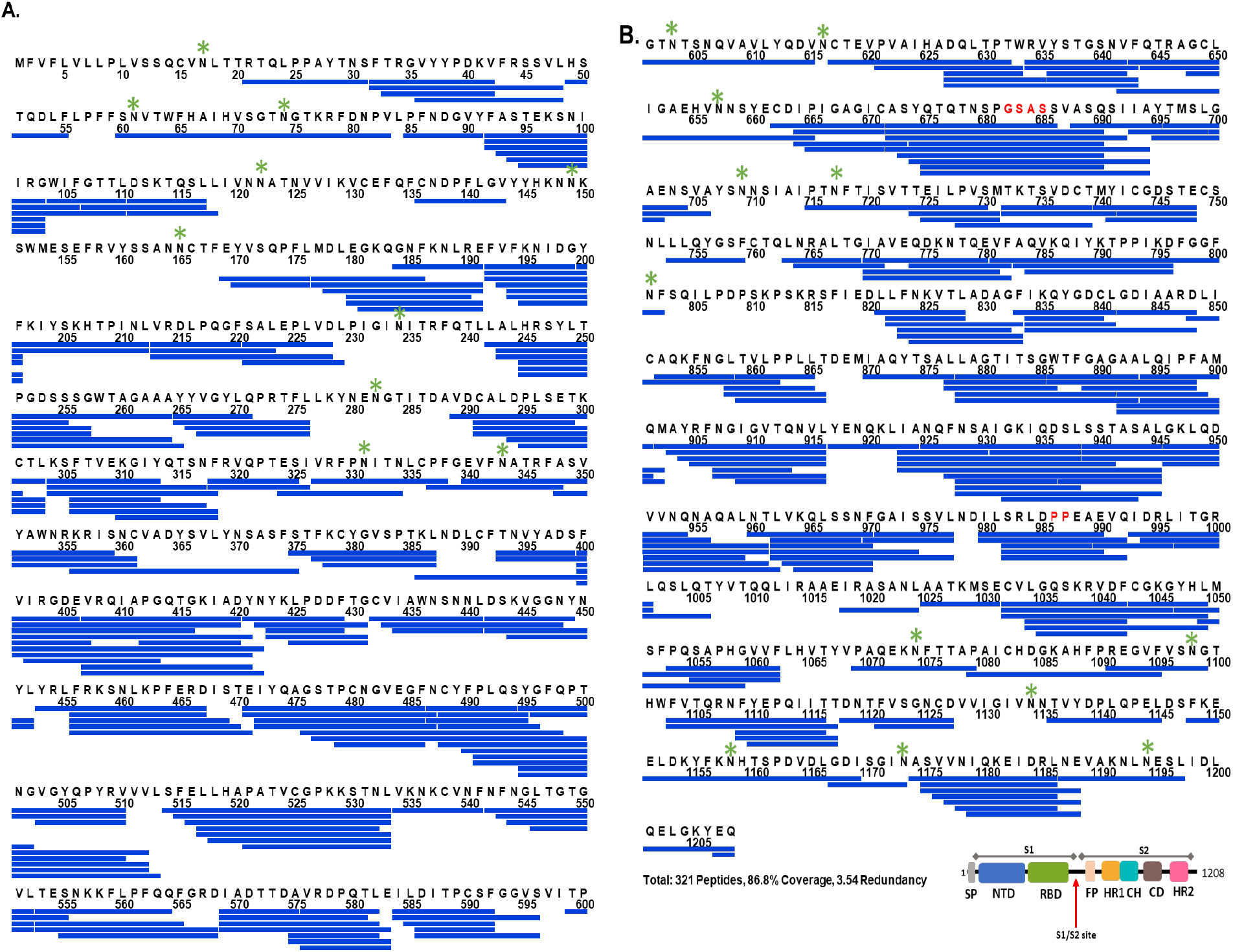
Primary sequence coverage map of pepsin proteolysed peptides of the S protein. Coverage map showing 321 peptides spanning 87% of the S protein: (A) 1 – 600 and (B) 601 – 1208, with the mutations highlighted in red. Glycosylation sites are indicated by asterisks (*) and peptide coverage for C-terminal twin strep-tag is not shown. The domain organization for S protein construct 1-1208 is shown.

**Figure S3:**
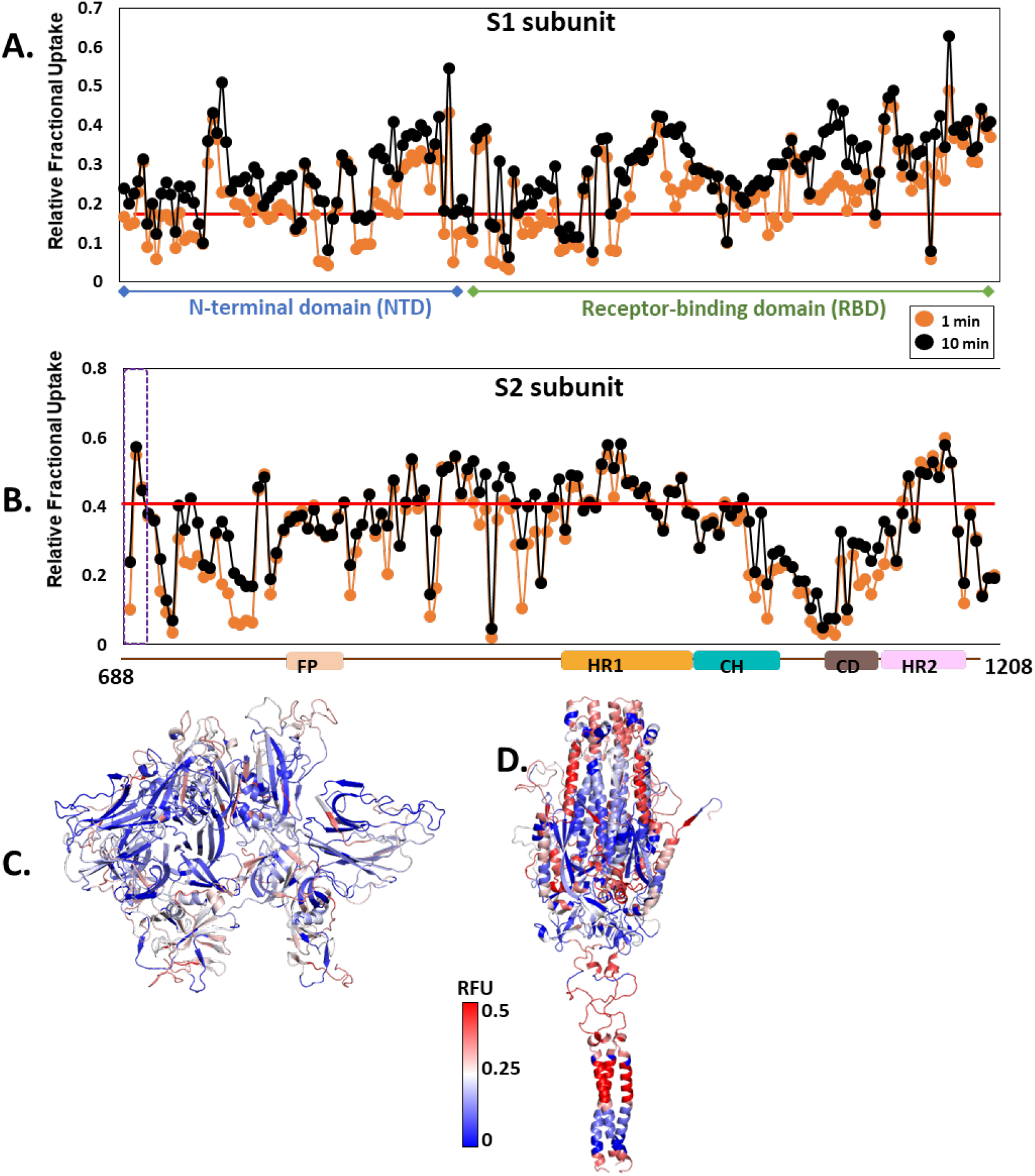
Time dependent changes in deuterium exchange for free S protein. Deuterium uptake of each pepsin proteolysed peptide listed from N-to C-terminus (X-axis) spanning S1 subunit (A) and S2 subunit (B) at deuterium labelling times 1 min and 10 min are represented as relative fractional uptake (RFU, Y-axis) values. Red line indicates the average RFU value. RFU values at 1 min of deuterium labelling time mapped on to the structures of the S1 (C) and S2 (D) subunits. High and low exchanging regions are as per key.

**Figure S4:**
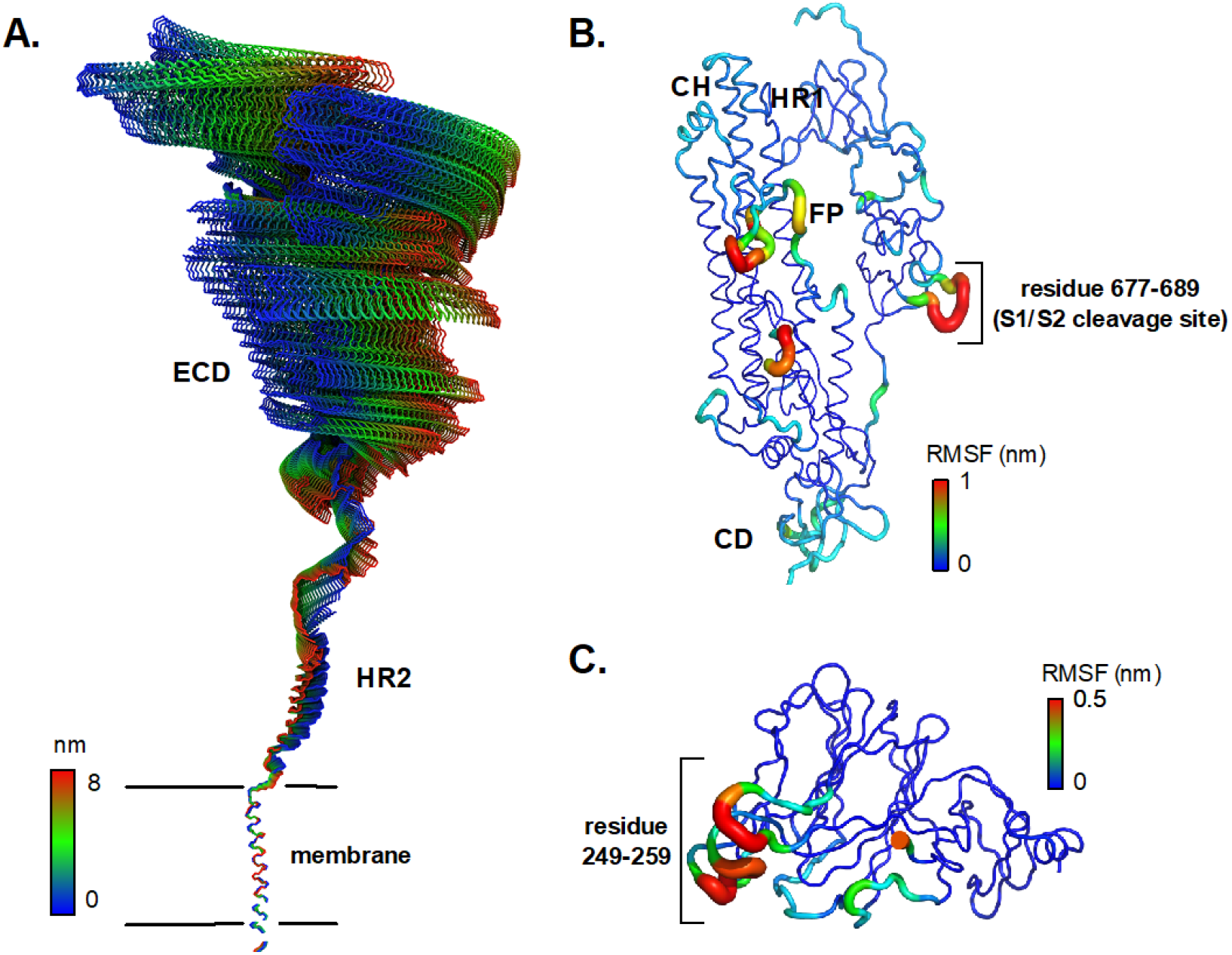
Dynamics of the S protein trimer from all-atom MD simulation. (A) The first principal motion of all backbone atoms for the full-length S protein during all-atom MD simulations as determined by principal component analysis (PCA). (B-C) RMSF values of backbone atoms on the S2 subunits and NTD. Residues with high RMSF are labelled.

**Figure S5:**
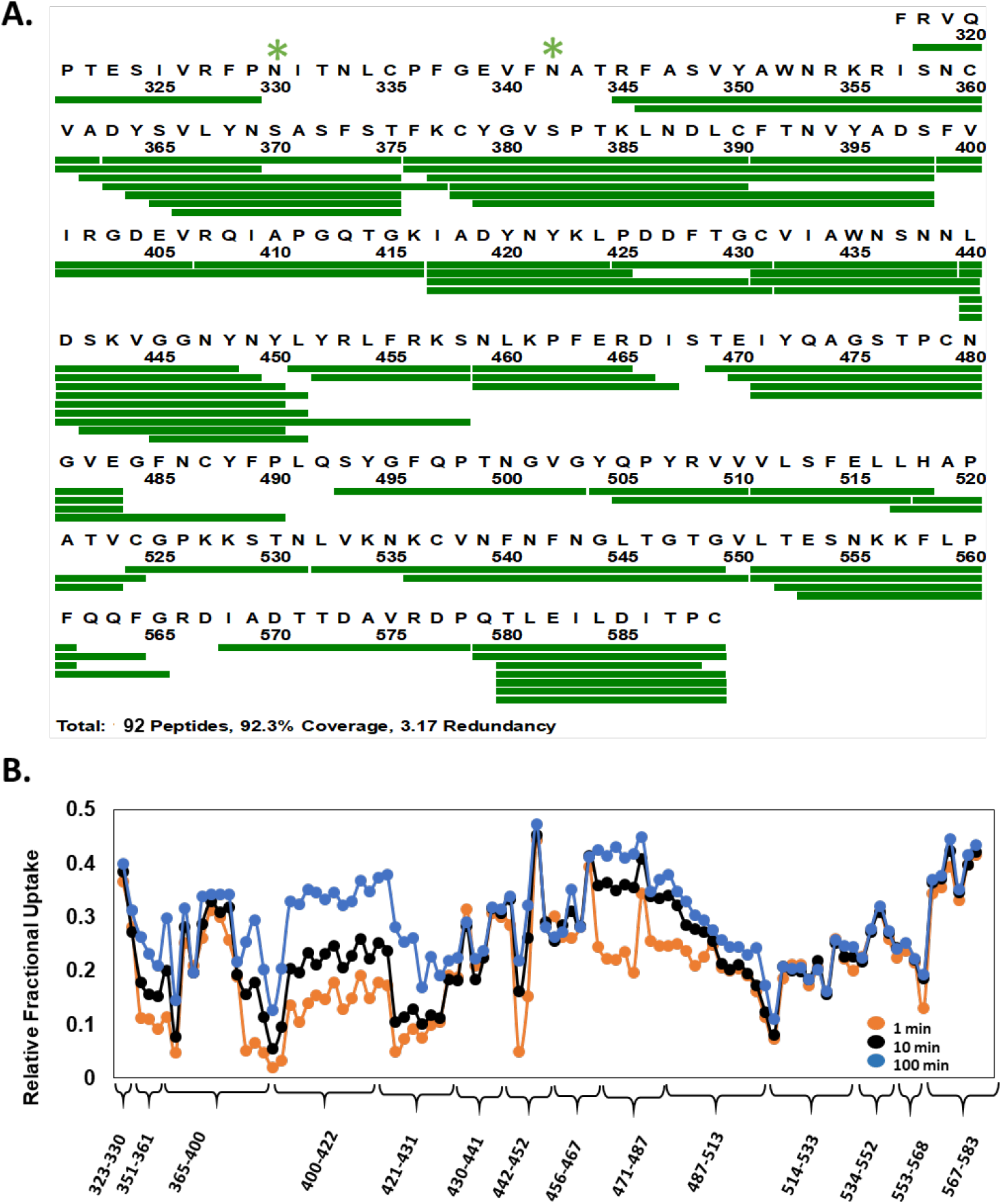
Primary sequence coverage and deuterium exchange profile of RBD_isolated_. (A) Coverage map showing 92 peptides (green bar) spanning ~92% sequence of MBP-RBD_isolated_ (318-589) fusion protein. N-terminal maltose-binding protein (MBP) affinity-tag is not shown. Glycosylation sites are indicated by green asterisk. (B) RFU plot of pepsin proteolyzed peptides of RBD_isolated_ listed N-to C-terminus for deuterium labelling times as per key.

**Figure S6:**
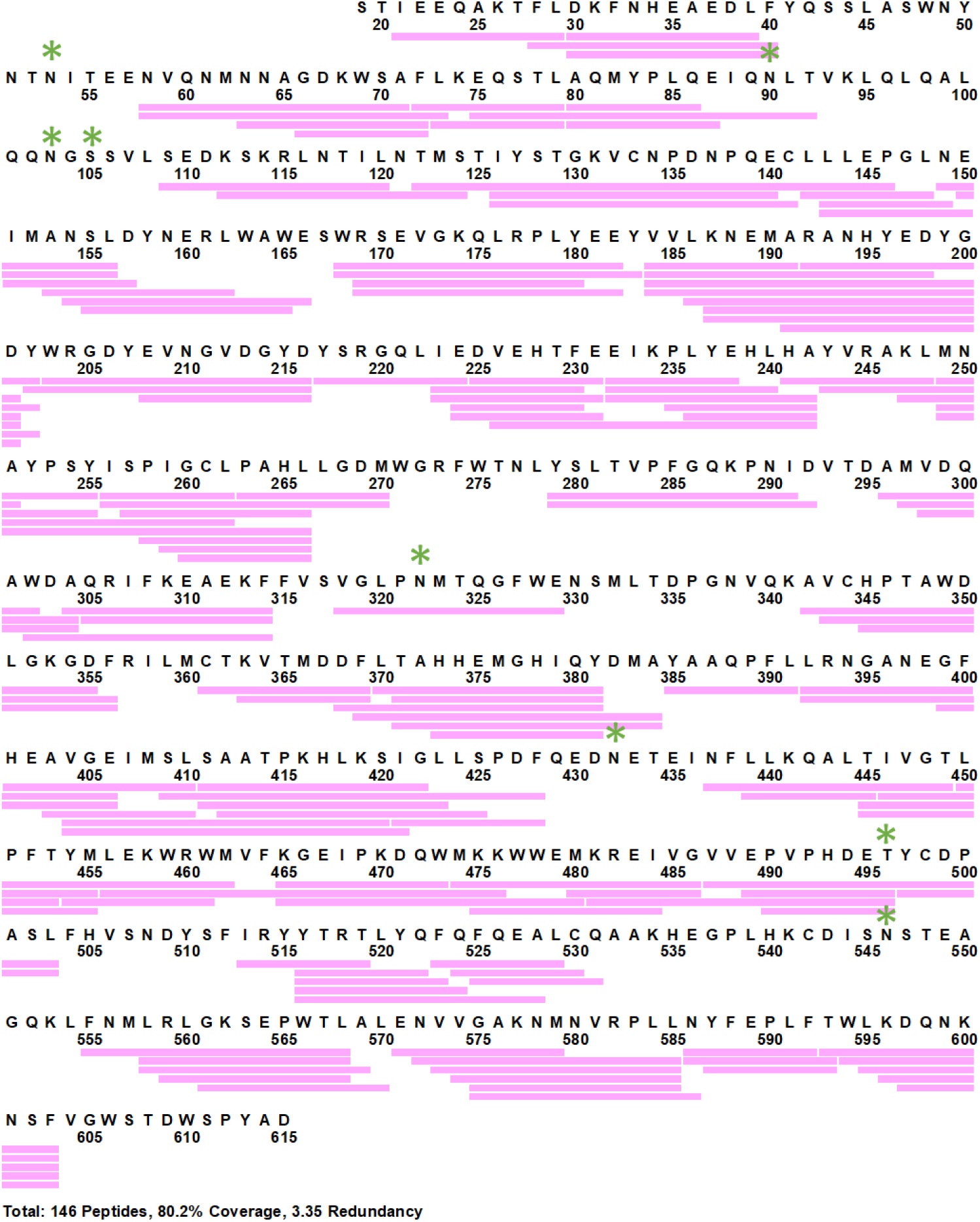
Pepsin digest map and sequence coverage ACE2. (A) Coverage map showing 140 peptides (pink horizontal bars) covering ~80% sequence of ACE2 (18-615). Sequence of FC-tag is not shown. Glycosylation sites are represented by green asterisk.

**Figure S7:**
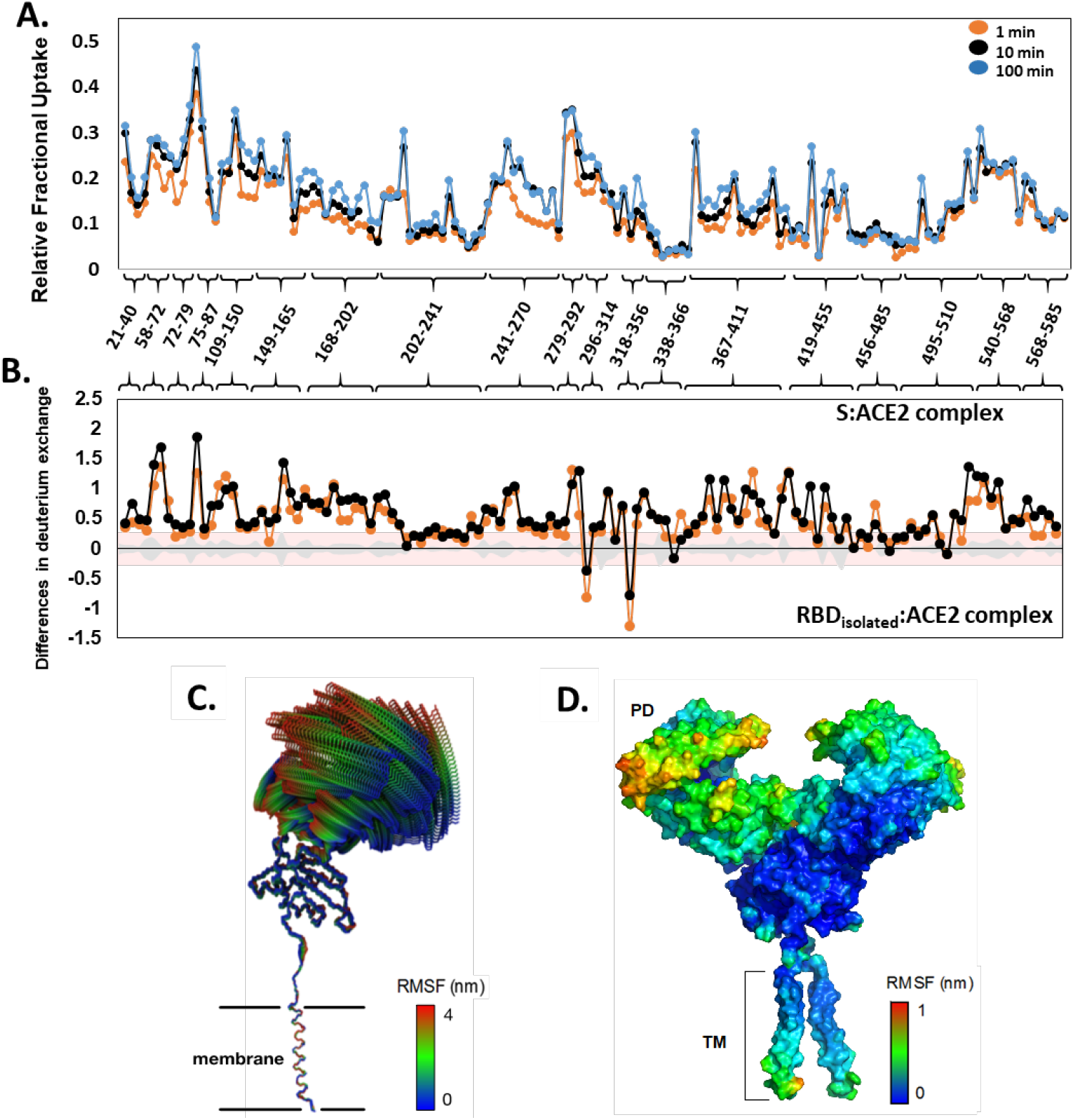
Deuterium uptake profile for ACE2 receptor and all-atom MD simulation of the ACE2-B0AT1 complex. (A) RFU values of pepsin proteolysed peptides listed in N-to C-terminus of ACE2 (peptide 18-615) for deuterium labelling times are shown. (B) Differences in deuterium exchange (Y-axis) of ACE2 peptides listed from N-to C-terminus (X-axis) between S:ACE2 complex and RBD_isolated_:ACE2 complex. Deuterium exchange significance threshold of ±0.3 D is indicated in red and standard errors in gray. (C) The first principal motion of the all backbone atoms of the ACE2 monomer as determined by PCA. (D) The RMSF values of the ACE2 receptor mapped onto the surface of the ACE2.

**Figure S8:**
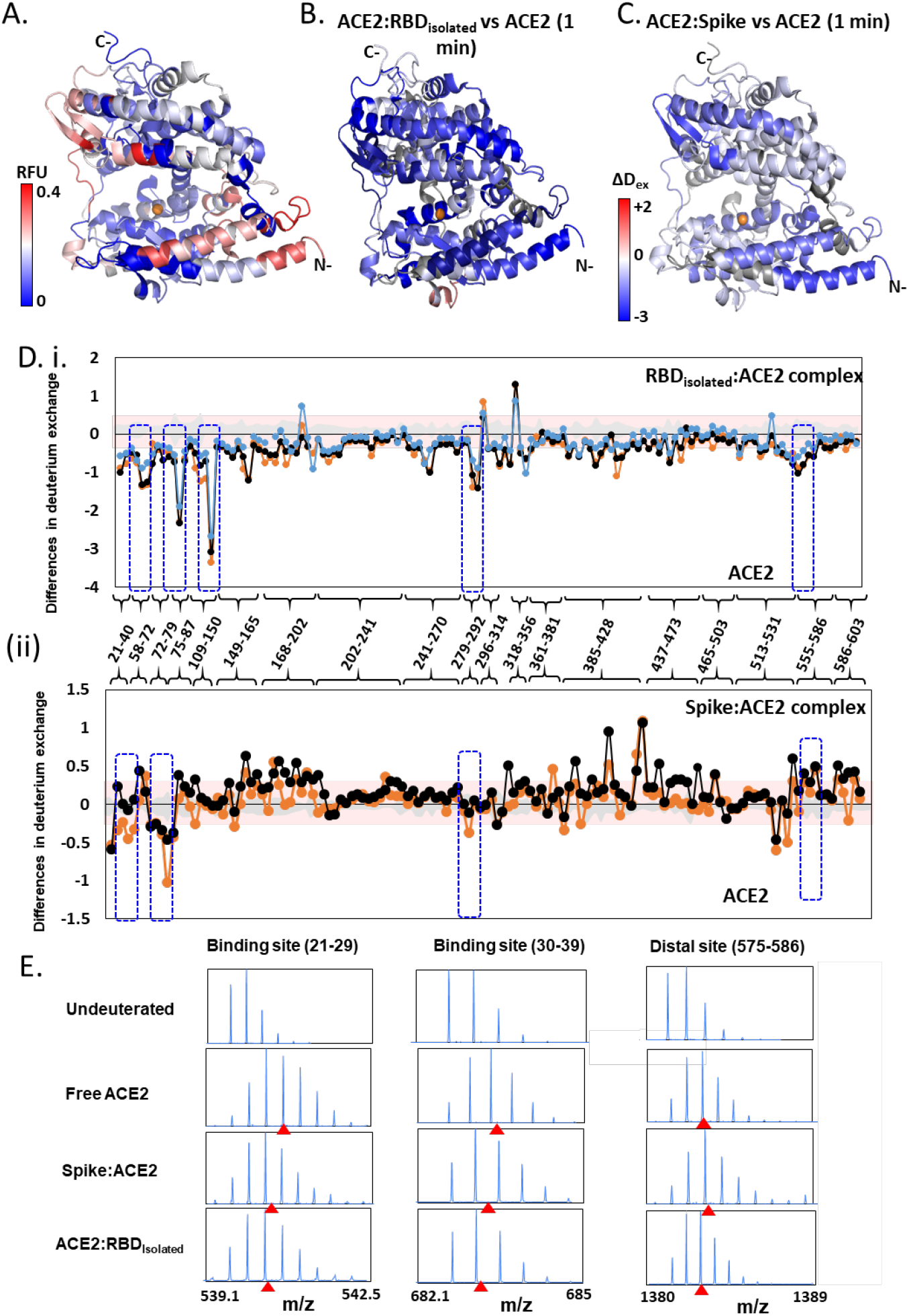
Effect of RBD_isolated_ and RBD_S_ complexes on ACE2 dynamics. (A) Structure of extracellular domain of ACE2 receptor (PDB ID: 1R42) depicting the RFU at t = 1 min. (B) Differences in deuterium exchange of RBD_isolated_:ACE2 complex and free ACE2 at t = 1 min is mapped onto the structure of ACE2, predominantly showing decreased deuterium exchange in ACE2 (shades of blue). (C) Heat map of differences in deuterium exchange (t = 1 min) of S:ACE2 complex and free ACE2. (D) Plot showing differences in deuterium exchange between ACE2 and complexes with RBD (i) and S (ii) at different labeling times. Pepsin digest fragments are indicated by their residue numbers. Cutoff ±0.3 D is the deuterium exchange significance threshold, indicated by pink shaded box, and standard errors are in gray. Positive differences denote increased deuterium exchange in (i) RBD:ACE2 or (ii) S:ACE2 compared to free ACE2, while negative differences denote decreased deuterium exchange. Peptides spanning the sites of interaction with RBD and two distal sites (278-292, 574-585) are highlighted. (E) Stacked mass spectra showing isotopic distribution for select peptides spanning the binding sites (21-29, 30-39) and a distal allosteric site (575-586) for ACE2, S:ACE2 and RBD_isolated_:ACE2 are shown at 1 min deuterium labeling time. Centroids indicated by red arrow-heads.

## Data S1 to S3

**Table S1:**

Relative Fraction uptake values at various deuterium labeling times for Spike and S:ACE2 complex

**Table S2:**

Relative Fractional Uptake values at various deuterium labeling times for free and ACE2-bound RBD (isolated)

**Table S3:**

Relative Fractional Uptake values at various deuterium labeling times for free ACE2 and its complexes with isolated RBD and Spike protein

